# Two distinct mechanisms for Na_v_1.7 null analgesia

**DOI:** 10.1101/2024.02.12.579826

**Authors:** Alexandros H. Kanellopoulos, Naxi Tian, James J. Cox, Jing Zhao, Christopher G. Woods, John N Wood

## Abstract

Genetic deletion and pharmacological inhibition are distinct approaches to unravelling pain mechanisms, identifying targets and developing new analgesics. Both approaches have been applied to the voltage-gated sodium channels Na_v_1.7 and Na_v_1.8. Genetic deletion of Na_v_1.8 in mice leads to a loss of pain, and antagonists are effective analgesics. Complete embryonic loss of Na_v_1.7 in humans or in mouse sensory neurons leads to profound analgesia substantially mediated by endogenous opioid signaling, and anosmia that is opioid independent. Autonomic function appears to be normal. Adult deletion of Na_v_1.7 in sensory neurons also leads to analgesia with diminished sensory neuron excitability but there is no opioid component of analgesia. Pharmacological inhibition of Na_v_1.7 leads to dramatic side-effects on the autonomic nervous system. Here we compare and contrast the distinct embryonic and adult null mechanisms of Nav1.7 loss-of-function analgesia. We describe an endogenous opioid mechanism of analgesia that provides new opportunities for therapeutic intervention and pain relief.

**Summary:** In contrast to Na_v_1.8, Na_v_1.7, a genetically validated human pain target is unsuitable for small molecule drug development because of its wide spread expression both centrally and peripherally.

## Introduction

The peripheral nervous system is the driver of conscious pain sensing in the brain. Inflammatory mediators and interactions with the immune system control pain thresholds and the gain of activity of damage-sensing neurons. These neurons release glutamate and neuropeptides within the dorsal horn of the spinal cord in response to tissue damage. Importantly, all analgesic drugs in clinical use also work well in mice, supporting this animal as a useful model system to explore pain mechanisms. The easy availability of DNA sequencing has allowed us to identify human pain genes ^1^. In principle, mechanistic studies of mouse models should provide us with a direct route to analgesic drugs, desperately needed in these days of the opioid overdose crisis.

Unfortunately, there is remarkable redundancy in signal transduction in the peripheral pain system. For example, there are at least four major heat sensors in mice and humans ^2^. Thus attacking nociceptor transduction is generally unappealing for analgesic drug development. Moving away from transduction, the ion channels involved in electrical transmission are less redundant and the sodium channels Nav1.7 and Nav1.8 have been unambiguously linked to human pain and damage sensing through detection of functional variants of the *SCN9A* and *SCN10A* genes ^1^. The encoded channels are the targets of the miraculous analgesic lidocaine, for which visitors to the dentist give thanks! Lidocaine was derived from a plant indole identified in studies of pest-resistant barley. Its long lasting effects as an analgesic and superiority to procaine and cocaine were established in human trials in 1944. Low dose systemic lidocaine is an effective pain treatment, but high doses can cause death. Thus Nav1.7 and Nav1.8 selective blockers are of considerable interest, especially as sensory neuron Nav1.7 and Nav1.8 null mice are healthy but have diminished pain sensing^1^.

## Results

### Nav1.8 as a pain target

The tetrodotoxin-resistant sodium channel Nav1.8, cloned in 1996 ^3^, is selectively expressed in sensory neurons and has been shown to play an important role particularly in inflammatory and mechanical pain ^4^. Antagonists are potent analgesics in preclinical models of neuropathic and inflammatory pain ^5^. Gene therapy with antisense oligonucleotides ^6^ or dead Cas9 inhibition of Nav1.8 expression produces analgesia (Figure 1), but these studies have been overtaken by orally active antagonists that work in humans ^8^.

**Figure 1.**
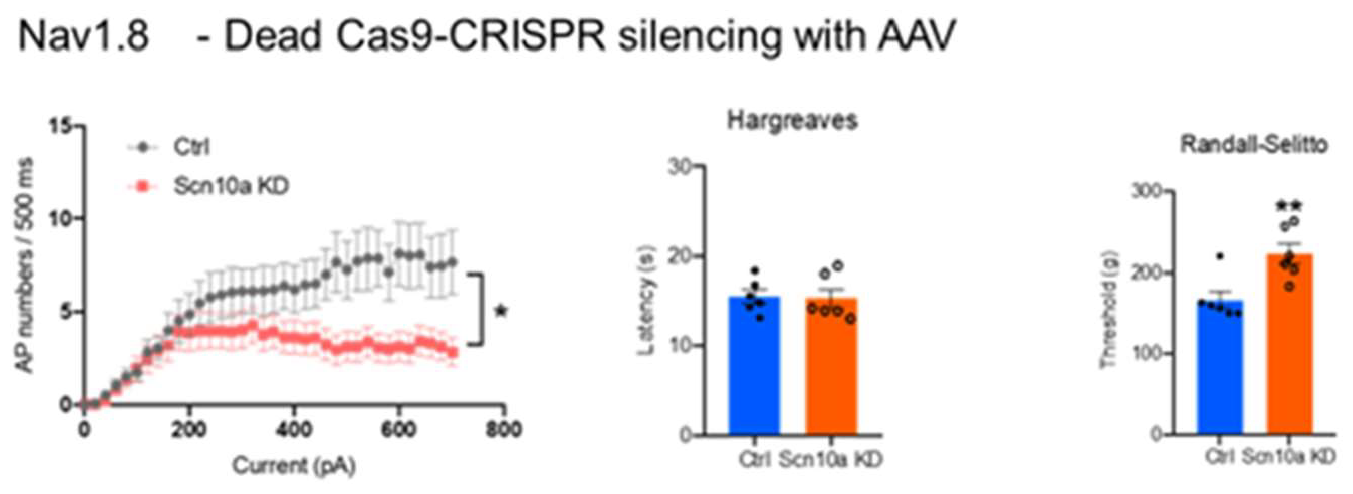
Epigenetic inhibition of Nav1.8 expression results in mechanical analgesia. AAV mediated delivery of targeted dead CMV-dSaCas9-ZIM3-pA results in lowered TTX-resistant sodium channel activity in sensory neurons with concomitant loss of noxious mechanosensation in behavioral assays. N=6 for behavioral assays n>10 for electrophysiological studies.

**Figure 2.**
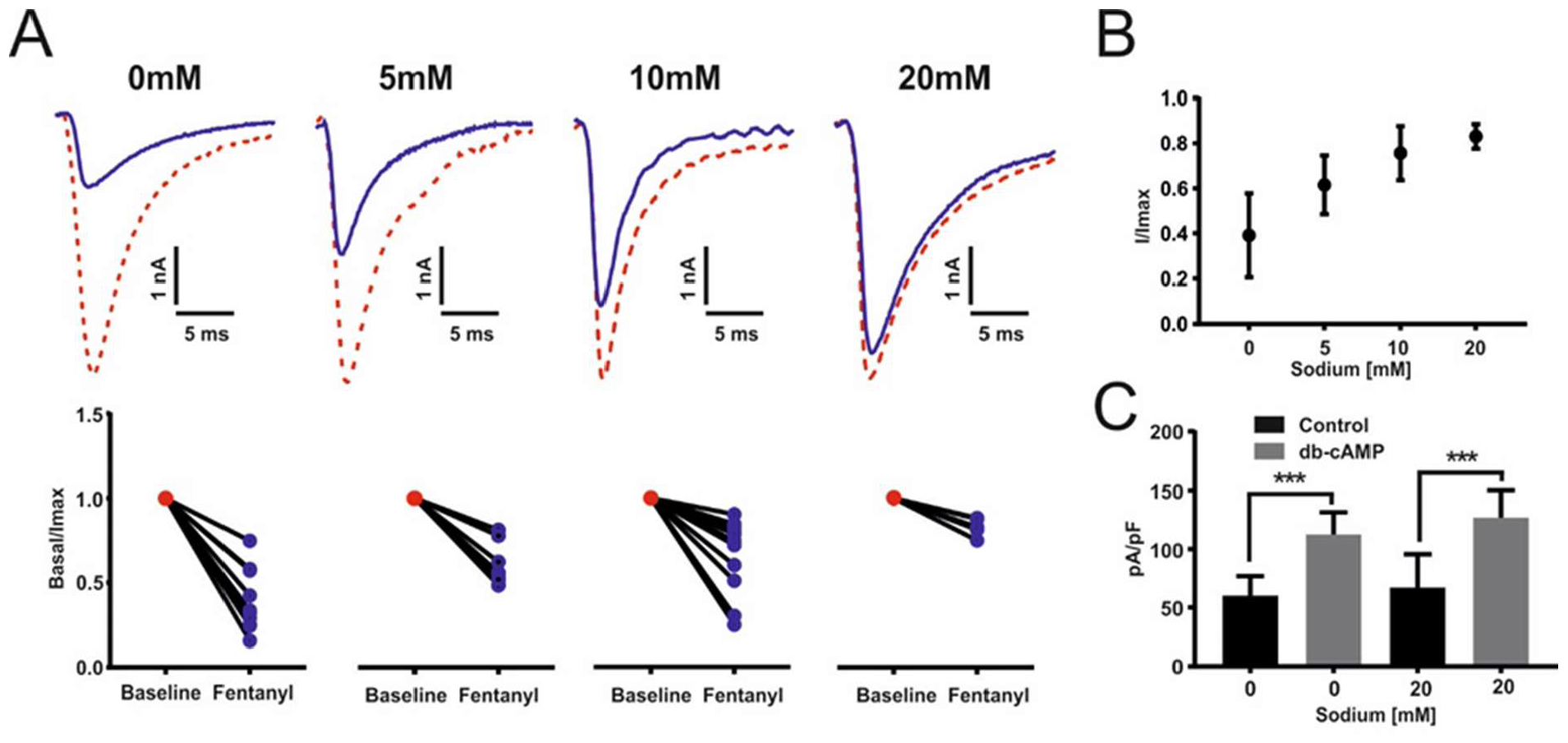
Effect of varying intracellular sodium on opioid inhibition of Nav1.8 currents. A) Electrophysiological example traces of TTXr Nav1.8 currents from dorsal root ganglia neurons following the exposure to 100nM fentanyl at different intracellular concentrations of sodium (0mM, 5mM, 10mM and 20mM). Corresponding dot plot of TTXr Nav1.8 current following the exposure to 100nM fentanyl at different intracellular concentrations of sodium 0mM n=14, 5mM n=10, 10mM n=14 and 20mM n=6. All currents are normalised and compared to baseline. Student’s *t* test, 0mM p<0.001, 5mM p<0.001, 10mM p<0.001 and 20mM p<0.05. (B) Data from (A) plotted as normalised peak currents compared to sodium concentration. Data represents ± SEM. (C) Electrophysiological recordings of TTXr Nav1.8 currents in dorsal root ganglia neurons following exposure to db-cAMP in 0mM and 20mM intracellular concentrations of sodium. WT 0mM vs db-cAMP 0mM, ***p<0.0002, change from baseline Δ=52.3pA/pF. WT 20mM vs db-cAMP 20mM, ***p<0.0006, change from baseline Δ=59.8pA/pF. No significant change between db-cAMP 0mM vs db-cAMP 20mM.

Many Na_v_1.8-focused drug development programs were halted after the discovery of a genetic link with Brugada sudden death syndrome and Na_v_1.8. This issue has been brilliantly resolved by Christoffels et al. who showed that a cryptic intronic promoter drives the production of a C-terminal fragment named SCN10A-*short* comprising the last 8 transmembrane segments of Na_v_1.8 in heart ^7^. There, this inactive protein promotes the activity of the heart channel Na_v_1.5, explaining why the loss of SCN10A-*short* can result in cardiac dysfunction and Brugada sudden death syndrome. This may explain the absence of human bi-allelic loss of function Na_v_1.8 mutants with diminished pain. The loss of Na_v_1.8 is likely to lead to cardiovascular dysfunction during development that is lethal. These insights demonstrate that cell line expression of human Na_v_1.8 and human Na_v_1.8-*short* plus Na_v_1.5 for drug screening will be able to identify candidate small molecules without cardiac effects for further development as human analgesics. An orally active Na_v_1.8 antagonist that is analgesic for acute pain has succeeded in phase 3 trials ^8^.

Recently phase 2 evidence of a useful effect on pain associated with diabetic neuropathy with a statistically significant and clinically meaningful reduction in the primary endpoint (clinical trial NCT05660538) was obtained. These encouraging data should stimulate others to take up the Na_v_1.8 challenge, particularly because no side effects were associated with the treatment.

### Na_v_1.7 as a pain target

A major reason for a change of focus from Na_v_1.8 to Na_v_1.7 was the remarkable demonstration that Na_v_1.7 was required for pain in both humans and mice ^1,9,10^. A genetic link with three human pain disorders was also found for dominant activating variants of this channel; a subset of severe Primary Erythermalgia cases, a similar but lesser phenotype in some cases of small fiber neuropathy, and the distinct phenotype Paroxysmal Extreme Pain disorder, (OMIM 133020 and 167400). Given the role of Na_v_1.7 in sympathetic neurons in heat pain in mice ^11^, it is possible that hyperactive Na_v_1.7 channels in sympathetic neurons play a significant role in the very rare cases of Primary Erythermalgia linked to Na_v_1.7 mutations (OMIM 133020). However, mouse models of gain-of-function human Na_v_1.7 mutants do not show an erythermalgia phenotype, whilst a human mutation linked to painful small fiber neuropathy does exhibit enhanced pain, interestingly in mice in a sex dependent manner ^12^. This fits with the observations of diminished C-fiber peripheral terminals in some human Na_v_1.7 pain free nulls ^13^. The inactivation-defective Na_v_1.7 mutant sensory neurons characteristic of paroxysmal extreme pain disorder (PEPD) linked to mechanical pain are more plausibly linked to mechanosensitive sensory neuron dysfunction ^14^. These findings, taken together, make Na_v_1.7 the best-validated human pain target to have been discovered.

### Embryonic Na_v_1.7 null opioid-dependent pain-free mice and humans

Embryonic deletion of Na_v_1.7 in sensory neurons leads to analgesia, but does not alter sensory neuron excitability in mice ^15^. As Na_v_1.7 plays an important role in sensory neuron action potential generation ^16^, this requires that compensatory effects rescue the excitability of sensory neurons in embryonic nulls. Na_v_1.7 is the principal human parasympathetic sodium channel, and plays an important role in sympathetic neurons as well as throughout the CNS ^17^ and potentially in insulin release ^18^. Therefore there must be compensatory upregulation of other voltage-gated sodium channels in the embryonic nulls to rescue central, autonomic and sensory function. Analysis of mRNA transcripts in Na_v_1.7 null sensory ganglia do not reveal enhanced transcription of other sodium channels ^19^, so compensation must occur at a later stage of functional protein expression such as trafficking. Sodium channels exist as dimers, and mass spectrometry of sensory neuron epitope-tagged Na_v_1.7 shows that B_2_ and B_3_ subunits, as well as Na_v_1.1 associate with Na_v_1.7 ^20^. Na_v_1.1 (13 hits 6.7% coverage on mass spectrometry of immunoprecipitates) is thus a candidate to replace Na_v_1.7 in peripheral neurons of embryonic null mice.

Although sensory neurons appear completely normal in mouse Na_v_1.7 nulls, they nonetheless show a dramatic loss of Substance P and glutamate release on depolarization. Interestingly synaptotagmins are physically associated with Na_v_1.7 ^20^. This inhibition of neurotransmitter release can be partially reversed by naloxone, as can analgesia in both mice and humans, suggesting that the opioid-dependent block of neurotransmitter release ^21^ plays a major role in Na_v_1.7 embryonic null analgesia. Importantly, the anosmia associated with loss of Na_v_1.7 seems to only depend on diminished electrical activity rather than opioid induction ^22^– a quite different mechanism from that which occurs in embryonic null somatosensory neurons. There is increased PENK mRNA expression in embryonic Na_v_1.7 null sensory neurons, but increasing enkephalin levels to a similar level by deleting transcription factor NFAT5 does not cause analgesia ^23^. As well as increased opioid peptide expression, the opioid signaling pathway in sensory neurons was found to be massively potentiated in embryonic Na_v_1.7 null mice by the Hucho group, and this seems to be the critical element in opioid signaling that leads to analgesia in Na_v_1.7 nulls ^24^.

An intimate association between Na_v_1.7 and μ-opioid receptors in the membrane is consistent with the role for this receptor demonstrated by Pereira et al^23^ in Na_v_1.7 null analgesia (see below) and the findings of the Hucho group ^24^. One of the G-proteins involved in μ-opioid signaling, GNAO1 is linked to Nav1.7 in studies of epitope tagged Na_v_1.7 interacting proteins, consistent with a close physical relationship between Na_v_1.7 and μ-opioid receptor expression ^25^

### Adult null Na_v_1.7 opioid-independent analgesia

Importantly, if Na_v_1.7 is deleted specifically in adult mouse sensory neurons with tamoxifen-inducible Cre recombinase, analgesia is also obtained ^16^. However, in these experiments in contrast to embryonic nulls, there is a dramatic loss of electrical excitability in sensory neurons and no apparent role detected for the opioid system This difference with the findings in embryonic nulls ^19, 15,25^ supports the view that there are two distinct types of analgesia associated with embryonic or adult loss of Nav1.7 expression. As Na_v_1.7 is the voltage-gated channel that responds first with action potentials to sensory neuron depolarization, the adult data make sense ^26^

This significant observation in no way argues against the findings of profound analgesia in embryonic null humans and mice reversed by naloxone, but raises important issues for drug development. Consistent with the broad expression, sometimes at low levels, of Nav1.7 in most neuronal sub-population in the CNS (82 out of 96 sets of central neurons in MouseBrain.org), as well as in the autonomic nervous system and non-neuronal tissues like the pancreas, side effects are the major problem with Na_v_1.7 antagonists as potential analgesics. Na_v_1.7 has a key physiological role throughout the CNS in synaptic integration, and is known to regulate weight in mice ^17^. Thus centrally acting blockers that diminish neurotransmitter release will have major side effects. Peripheral autonomic side effects are also dramatic and have been found in adult mouse studies of Nav1.7 antagonists^16^. Side effects thus rule out Nav1.7 as a target for small molecule analgesic drug development. Of course, these side effects also impact on other potential applications for Nav1.7 antagonists. A report of a role in arthritis of Nav1.7 in chondrocytes relies on deletion by a chondrocyte-specific Cre recombinase, (although this Cre is also expressed in hind brain and other neurons that express Na_v_1.7 (see MouseBrain.org)). The same side effect issues that rule out analgesic development also apply to this potential therapeutic target for osteoarthritis ^27^.

### Nav1.7 is not a small molecule analgesic drug target

It is clear that small molecule Na_v_1.7 antagonists will always be bedeviled with side effects due to the broad expression of Na_v_1.7 in the pancreas, autonomic nervous system and throughout the brain ^17^. The reason that such side effects are not apparent in embryonic Na_v_1.7 nulls must be a consequence of compensatory expression of other sodium channels that rescue the excitability of autonomic neurons, CNS neurons throughout the brain (see MouseBrain.org) and sensory neurons. Genetic data from embryonic nulls are thus no basis for deciding that Na_v_1.7-targeted drugs will be side-effect free. In fact, major side effects have been observed with potent Na_v_1.7 antagonists ^16^. For these reasons, Na_v_1.7 is not a viable small molecule analgesic target, but the embryonic null data provide us with an exciting new insight into a potential analgesic strategy exploiting altered opioid signaling.

### Endogenous opioid mediated analgesia

How does activation of the opioid system occur in embryonic nulls? Na_v_1.7 carries a persistent sodium current and is expressed near opioid receptors involved in analgesia. Immunoprecipitation of TAP-tagged Na_v_1.7 shows the presence of GNAO1, a μ-opioid receptor binding partner. Class-A G-protein coupled receptors such as opioid receptors have a transmembrane sodium binding pocket that, when occupied, inhibits downstream signaling. Is it possible that sodium ingress through Nav1.7 acts as a second messenger? In the presence of Na_v_1.7, sodium normally inhibits opioid GPCR regulation of neurotransmitter release. With the loss of Na_v_1,7, enhanced opioid signaling could occur if sodium block of GPCR activity was reduced ^28^. Intriguingly both PENK and NFAT5 mRNA expression are enhanced by lowering intracellular sodium ^23^. Of course sodium as a second messenger controlling potassium currents is a well established phenomenon, and a broader role is likely ^29^.

We examined the effect of changing intracellular sodium concentrations on the activity of μ-opioid receptors in individual mouse sensory neurons using a sensitive electrophysiological assay. Protein Kinase A (PKA) is known to phosphorylate five serine residues in the first intracellular loop of Na_v_1.8, a tetrodotoxin-insensitive voltage-gated sodium channel that is uniquely expressed in sensory neurons (*9)*. This results in a large increase in TTX-resistant sodium channel activity that can be quantitated by electrophysiological recording. Fentanyl, acting through μ-opioid receptors and Gi proteins can suppress the activity of PKA, and diminish the level of tetrodotoxin-resistant current (9,*10,11*). These observations provide us with a simple assay system for measuring opioid action in intact cells, that allows us to vary the level of intracellular sodium and examine the consequences.

We tested this concept by examining the functional expression of the TTXr sodium channel Nav1.8 and effects of external opioids with altered intracellular sodium (Figure 3,4). Here it can be seen that the inhibition of Na_v_1.8 functional expression by fentanyl can be potentiated in conditions of low sodium within sensory neurons in culture. The activation of PKA by dbcAMP is independent of altered sodium concentrations, suggesting that the opioid signaling mechanism itself is regulated by sodium. The proposed mechanism thus appears to be feasible. This hypothesis suggests that partial loss of Na_v_1.7 activity effected by drugs may not lower the levels of intracellular sodium to an adequate level to activate the opioid system to the level found with embryonic gene deletion. Pharma have generated a number of selective Na_v_1.7 antagonists that have weak analgesic action compared to the effects of gene deletion. Such drugs are also bedeviled with autonomic side effects^16^ whilst some effective but less selective compounds e.g. CNV1014802 are lidocaine-like and non-specific ^30^.

**Figure 3.**
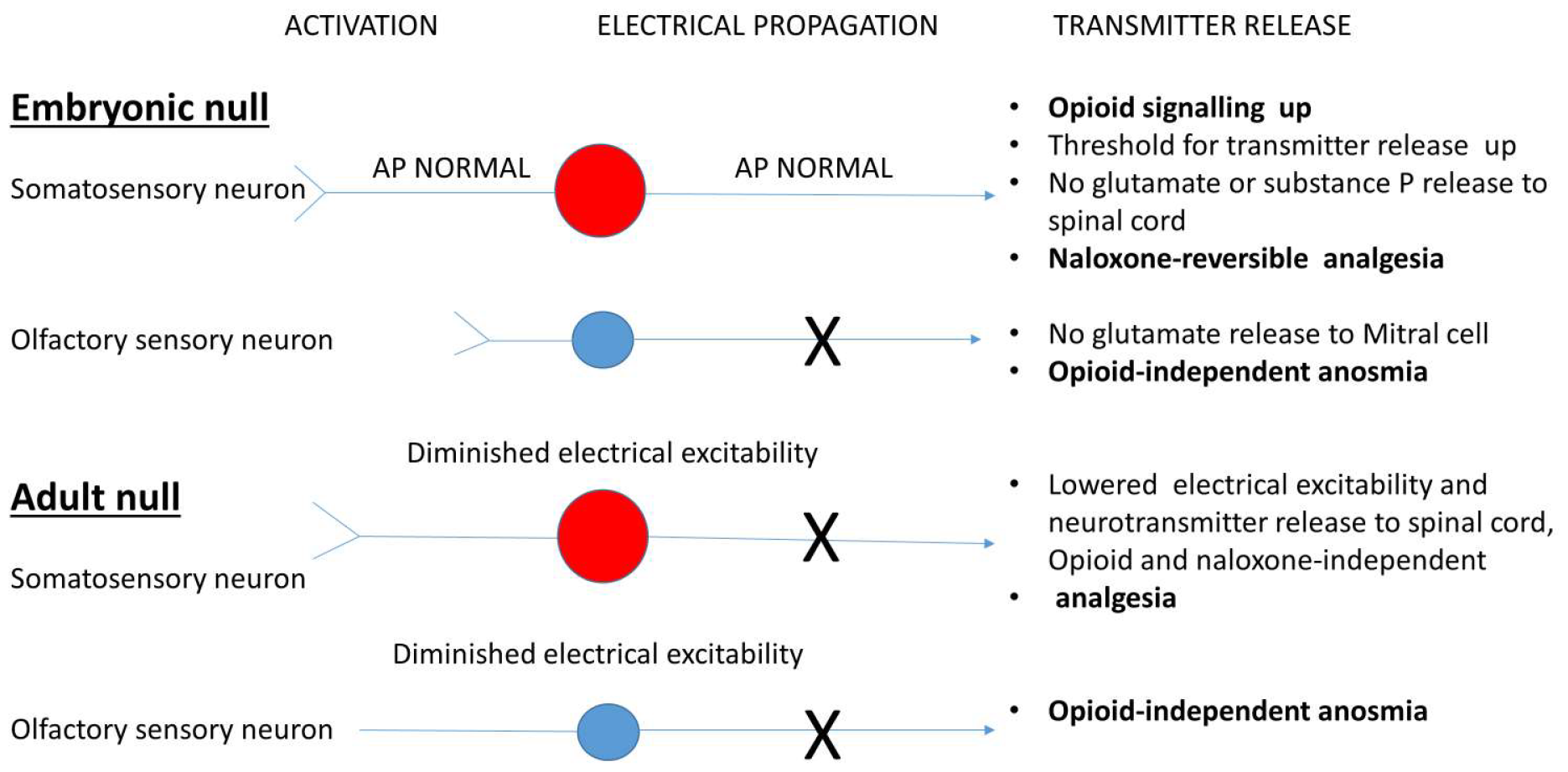
Distinct embryonic and adult mechanisms of analgesia associated with loss of expression of Na_v_1.7. AP = action potential. See references 15, 16,19, 22.

## Discussion

### Comparing Nav1.7 null analgesic mechanisms

Importantly, if we step away from the focus on Na_v_1.7 electrical activity, the induction of the opioid system in embryonic null sensory neurons suggests a new route to develop analgesia (see Figure 3). Evidence suggests that lowering intracellular sodium levels can activate the endogenous opioid system. Human embryonic Nav1.7 nulls have a lifetime of endogenous opioid-induced analgesia without respiratory depression, euphoria or constipation (although they all have Charcot’s joints by teenagehood and other risks inherent on having no pain to guide their behaviours). The opioid system is turbocharged in Nav1.7 null sensory neurons ^23,24^. Does the induction of the opioid system require complete (genetic deletion) rather than partial (pharmacological) loss of Nav1.7 activity to induce opioid-mediated analgesia via lowered sodium levels? Expensive sensory neuron specific gene therapy (ASOs, siRNA, CRISPR) that profoundly lowers Nav1.7 levels only in sensory neurons may be considered as a potential route to analgesia, but Nav1.7 in sympathetic neurons is clearly also important for pain.

Defining the precise mechanisms through which embryonic Nav1.7 channel loss leads to opioid signaling potentiation, neurotransmitter release loss and profound analgesia promises to be the most useful translational aspect of Nav1.7-related pain studies. A first step will be to test the hypothesis of a role for altered intracellular sodium in controlling opioid activity and transmitter release within sensory neuron terminals

Embryonic sensory neurons from Na_v_1.7 null mice are apparently normal in all respects. However, release of substance P and glutamate is severely compromised. In contrast embryonic null olfactory neurons are electrically silent. Adult nulls show some analgesia and anosmia linked to lowered excitability of sensory neurons and olfactory neurons respectively. The opioid signaling system plays no role in adult null Na_v_1.7 pain or olfactory deficits.

## Conclusions

The story of Na_v_1.7-focused analgesic development raises some interesting issues. Firstly, mechanistic studies of the events linked to Na_v_1.7 loss *in utero* and in adults have clearly been important in terms of drug development strategies, where misinformation has played a significant role in Pharma failure ^31^. Secondly, mice and humans seem to be very similar in terms of mechanisms. Thirdly, with Na_v_1.8 antagonists on the horizon, the future for pain treatment is looking brighter than for some time. It is ironic that genetic studies of an embryonic deletion of the pain related Na_v_1.7 sodium channel in humans and mice should lead us back to the sensory neuron opioid system, the most wonderful endogenous analgesic system that, could we manipulate it effectively, would save much human suffering. Finally, many of these studies have built on classic insights from Jessell, Iversen, Pasternak, Pert and Snyder. With the historically well-known role of a tetrodotoxin-resistant sodium channel in pain, it is very much “back to the future” for contemporary pain research.

## Materials and Methods

### Animals

All animal experiments were approved by the United Kingdom Home Office Animals Scientific Procedures Act 1986. Experiments were conducted using both female and male mice. For experiments using transgenic mice, wild-type littermate animals were used as controls. All strains of mice used for procedures were of C57Bl/6 background. All mice used for experimentation were at least 6 weeks old.

### Adeno-associated viruses

Three different AAV plasmids were assembled using Gibson assembly and sent for packaging into AAV9 (VectorBuilder): (1) CMV-dSaCas9-ZIM3-pA; (2) U6-sgRNA A, U6-sgRNA B, CMV-tdTomato-WPRE-pA, U6-sgRNA C, U6-sgRNA D; and (3) CMV-tdTomato-WPRE-pA. Guide RNA sequences were designed to map to the promoter of *Scn10a* (sgRNA A: GCCCGTCCTTAGCAGGATGGG; sgRNA B: GGGGGACAAAACACGCTTTG; sgRNA C: CTACAAGGAATCACGCCTTC and sgRNA D: GGCGTGATTCCTTGTAGATCC). C57BL6/J (9 weeks old, n=6 per group) were injected by the retro-orbital route with either AAVs 1 and 3 (control mice) or AAVs 1 and 2 (test mice). Pain behavioural tests were carried out from 14 days post injection.

### Electrophysiology

All electrophysiological recordings were performed using an Axopatch 200B amplifier and a Digidata 1440A digitizer (Axon Instruments), controlled by Clampex software (version 10, Molecular Devices). Filamented borosilicate microelectrodes (GC150TF-7.5, Harvard Apparatus) were coated with beeswax and fire-polished using a microforge (Narishige) to give resistances of 2 to 3 megohms. For voltage-clamp experiments, the following solutions were used. The extracellular solution contains 70 mM NaCl, 70 mM choline chloride, 3 mM KCl, 1 mM MgCl2, 1 mM CaCl2, 20 mM tetraethylammonium chloride, 0.1 mM CdCl2, 300 nM TTX, 10 mM Hepes, and 10 mM glucose (pH 7.3) with NaOH. The intracellular solution contains 140 mM CsF, 1 mM EGTA, 10 mM NaCl, and 10 mM Hepes (pH 7.3) with CsOH. Unless otherwise stated, standard whole-cell currents were acquired at 25 kHz and filtered at 10 kHz (low-pass Bessel filter). After achieving whole-cell configuration, the cell was left for 5 min to dialyze the intracellular solution. A holding potential of −100 mV was applied, and series resistance was compensated by ≥70%. All currents were leak-subtracted using a p/4 protocol. To record TTXr sodium currents, we applied a depolarizing voltage-pulse protocol to the cell; the cell was held at −100 mV and then stepped to −15 mV for 50 ms before returning back to −100 mV. This step was applied every 5 s for the duration of the experiment. The cells were continuously perfused using a gravity-fed perfusion system. All electrophysiological data were extracted using Clampfit (version 10, Molecular Devices) and analyzed using GraphPad Prism software (version 6, GraphPad).

### Statistical Analysis

Statistical analysis were performed with either Student’s *t* test with respective post hoc tests. *P* < 0.05 was considered statistically significant. Voltage-clamp experiments were analysed using cCLAMP software and Origin (OriginLab Corp., Northampton, MA) software programs. Current density-voltage (pA/pF) analysis by measuring peak currents at different applied voltage steps and normalised to cell capacitance. Voltage dependent activation data was fitted to a Boltzman equation y = (A2 + (A1 − A2)/(1 + exp((Vh − x)/k))) * (x − Vrev), where A1 is the maximal amplitude, Vh is the potential of half-maximal activation, x is the clamped membrane potential, Vrev is the reversal potential, and k is a constant. All Boltzmann equations were fitted using ORIGIN software. ±SEM data were assumed to be normally distributed. Unpaired Student’s t test was used for statistical comparisons. Significance was determined at p < 0.05. Individual p values are given for each comparison made. Fentanyl dose response curve and IC50 calculations were fitted and measured using ORIGIN software.

## Acknowledgments

We thank members of our research teams for their input and tolerance. Our work was supported by the Wellcome Trust, Versus Arthritis, Cambridge NIHR BRC, and the MRC.

